# Python-based automation of INDIGO webserver using Selenium: A high throughput analysis of Sanger sequence data to detect allelic variations created by CRISPR/Cas9-mediated genome editing of crop plants

**DOI:** 10.1101/2025.08.04.668409

**Authors:** Vandana Suresh, Chandra Girish, V. S. Sresty Tavva

## Abstract

CRISPR-Cas9 has revolutionized plant genome editing by enabling precise introduction of insertion/deletion (indel) mutations, critical for functional genomics and crop improvement studies. Sanger sequencing, combined with bioinformatics tools like the INDIGO webserver from Gear Genomics, is essential for validating these mutations. However, manual analysis of large numbers of Sanger sequencing *(.ab1)* files is labor-intensive, particularly when analyzing multiple guide RNA (gRNA) target regions. We developed a Python-based automation pipeline using Selenium with integrated highlighting of protospacer adjacent motif (PAM) regions in the resulting HTML reports. This pipeline enhances scalability of Sanger sequence data analysis and improves result interpretability by automating PAM region identification and supporting multiple gRNA regions. This tool significantly accelerates CRISPR-Cas9-mediated mutation analysis, offering a high throughput, reproducible solution for genome editing research.

## Introduction

CRISPR-Cas9 has emerged as a transformative tool for precise gene editing in plants, enabling targeted modifications to enhance traits such as yield improvement, disease resistance, and environmental adaptability. This system utilizes a guide RNA (gRNA) to direct the Cas9 nuclease to a specific genomic locus, where it induces double-strand breaks (DSBs). These breaks are repaired by cellular mechanisms, primarily non-homologous end joining (NHEJ) or homology-directed repair (HDR), leading to insertions, deletions (indels), or precise sequence replacements (Jinek *et al*., 2012). In plants, CRISPR-Cas9 is widely applied to edit genes associated with agronomic traits, making it a cornerstone of modern crop improvement (Ricroch *et al*., 2017).

Confirmation of CRISPR-Cas9-induced edits is critical to validate the precision and efficiency of modifications. Two primary sequencing methods to confirm CRISPR edited lines are Deep sequencing (NGS) and Sanger sequencing. NGS provides high-throughput, comprehensive analysis of edited genomic regions (Feng *et al*., 2014). However, the high cost of NGS makes it less feasible for routine analysis of large populations of edited plant lines. Due to the high cost and resource demands of NGS, Sanger sequencing is often preferred for large-scale screening of edited plant lines.

Sanger sequencing is a cost-effective, reliable, and widely accessible method for confirming indels in CRISPR-Cas9 workflows. This method is used to sequence individual PCR-amplified target regions, producing high-quality, reliable data for detecting indels and base substitutions (Sanger *et al*., 1977). Sanger sequencing of edited plant lines, followed by analysis of the resulting.ab1 files, is a standard method which allows researchers to verify the zygosity (heterozygous or homozygous) of indels, facilitating the selection of desired mutant lines for further propagation and phenotypic evaluation (Yang *et al*., 2017).

By combining Sanger sequencing with bioinformatics platforms, researchers can accurately characterize CRISPR-Cas9 induced modifications. Among numerous bioinformatics tools available for Sanger sequencing data analysis, INDIGO webserver from Gear Genomics, developed by the European Molecular Biology Laboratory (EMBL, Heidelberg, Germany), stands out as an open-access, user-friendly platform specifically designed for analyzing CRISPR-Cas9-induced gene edits. INDIGO aligns sequencing reads against a reference sequence, separating alleles into high-quality alignments (Alignment 1 and Alignment 2) and visualizes the results. The tool deconvolutes chromatogram files (.ab1), enabling researchers to detect single nucleotide variants (SNVs) and insertions/deletions (indels) with ease.

INDIGO accepts inputs such as a chromatogram files of sample and reference files (.ab1 file) or short FASTA sequence and produces a detailed output page comprising several components: a standard trace viewer, alignments of the two identified alleles, relative to the reference, an alignment comparing the first allele to the second with estimated allelic fractions, a variant table that connects reference positions, base call positions, and signal trace positions through hyperlinks to the original trace, and a decomposition chart illustrating the error associated with each potential heterozygous indel length (Rausch *et al*., 2020).

INDIGO webserver can be used impeccably to analyze single sequence reactions at a time; this slows down handling of large datasets that will be generated as part of the product pipeline. Manual analysis of large number of sample files is time-consuming, prone to human error, and hindered by the challenge of instantly identifying the Protospacer Adjacent Motif (PAM) region, which is essential for validating CRISPR-Cas9 edits. To overcome these limitations and enable high-throughput analysis, we developed a Selenium-based automation script in Python for the INDIGO webserver.

In this article, we focus on the power of automation developed for data processing and interpretation for Indel mutational studies using INDIGO webserver. It facilitates batch processing of large sample sets, making it ideal for extensive analysis of genome edited lines. Additionally, the automation includes functionality to identify and extract PAM containing sequences, enhancing the efficiency of edit validation. This Python-based Selenium workflow transforms INDIGO into a high-throughput application, improving scalability and reliability in detecting mutations from large number of sequencing files (.ab1) generated from CRISPR-Cas9-mediated genome edited lines.

## Methods

### Overview of INDIGO Automation Pipeline

To enhance the throughput of Sanger sequencing data analysis, an automation pipeline was developed in Python (version 3.8) to process sequencing files (.ab1) using Selenium (https://www.selenium.dev/documentation/) for interaction with the GEAR Genomics INDIGO web tool (https://www.gear-genomics.com/indigo/). The pipeline automates the process of uploading.ab1 files containing sequencing data from CRISPR-Cas9 edited samples, along with a wild-type control sequence as a reference. It then submits these files for analysis and retrieves the resulting HTML report. The.ab1 files generated from genome edited lines were organized in a designated input folder, while the reference.ab1 file was stored in a separate wild-type folder for comparison. The Protospacer Adjacent Motif (PAM) sequence region, critical for validating CRISPR-Cas9 edits, was annotated using SnapGene Viewer (GSL Biotech LLC, Boston, MA, USA), which facilitated visualization and analysis of the target DNA sequence to confirm the PAM region relevant to the CRISPR experiment.

### Error Handling and Validation

The script included basic error handling to detect missing input folders and web element failures. If the input folder is not found, the script terminates with an error message. For each file, exceptions during processing (e.g., timeouts or missing elements) were caught and reported, ensuring robustness across multiple files.

This automated pipeline enabled efficient processing of sequencing data, with the highlighted PAM sequence region facilitating visual inspection of CRISPR editing outcomes in the resulting HTML files.

### System Requirements

The pipeline was developed and tested on a Windows 10 system with 16 GB RAM and an Intel^®^ Core^™^ i5 processor. The Chrome browser (version 126.0) and corresponding ChromeDriver were used for web automation. The input folder contained.ab1 files generated from Sanger sequencing of CRISPR-Cas9-edited rice lines, with a wild-type.ab1 file used as a reference.

## Results

The automation pipeline for analyzing Sanger sequencing (.ab1) files through the INDIGO web application follows a clear, step-by-step process, as shown in Figure 1. This flowchart highlights the key stages of automating CRISPR-Cas9 mutation analysis, ensuring an efficient and scalable approach. This workflow shows how the pipeline automates repetitive tasks, manages errors effectively, and integrates tools like Selenium and ChromeDriver with INDIGO, streamlining CRISPR-Cas9 mutation analysis for plant genome editing research.

**Fig. 1:**
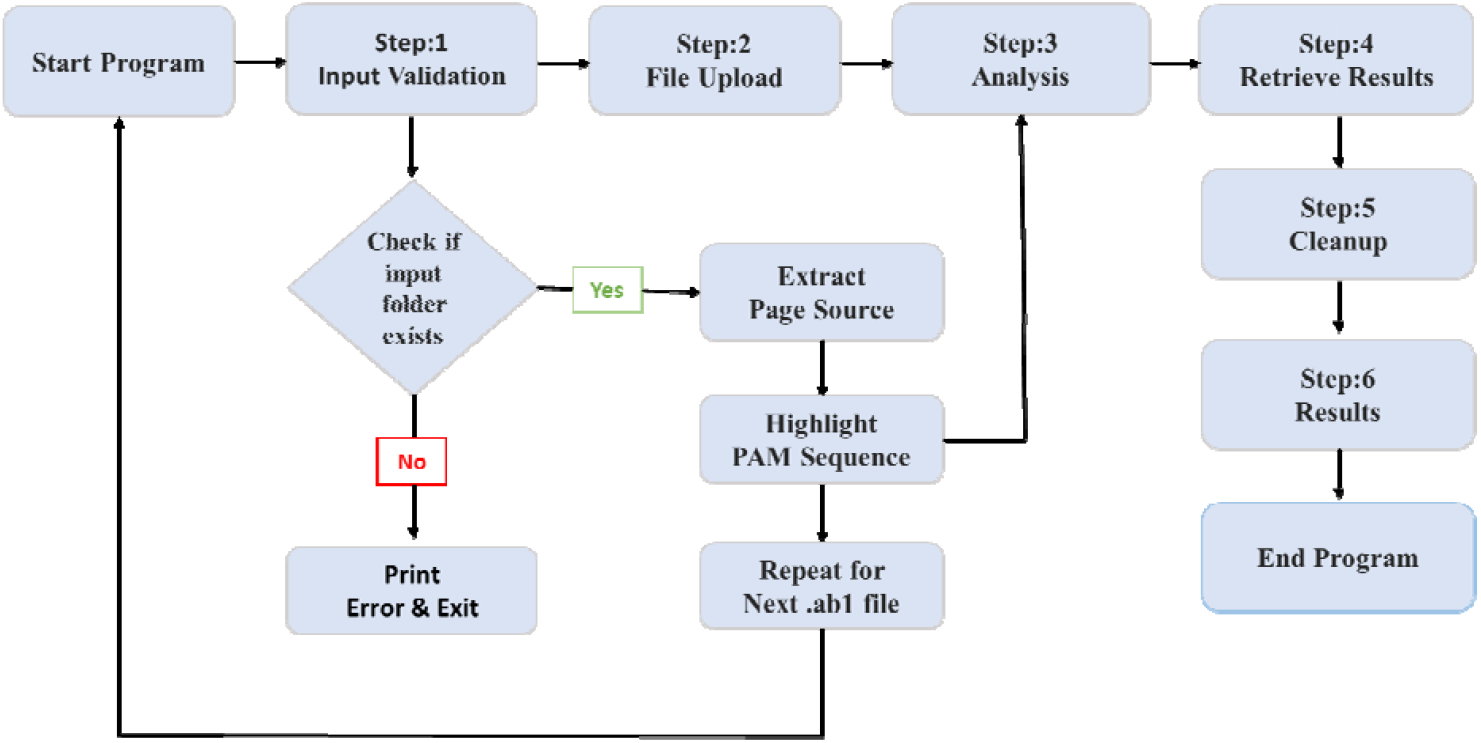
Workflow of the Python-based automation of INDIGO webserver using Selenium. The pipeline starts with the “Start Program” step, which launches the automation process. In **Step 1: Input Validation**, the pipeline checks if the folder containing the (.ab1) files exist. If the folder is missing, the pipeline displays an error message (“Print Error”) and stops (“Print Error & Exit”). If the folder is found, the workflow proceeds to the next step. **Step 2: File Upload**, where the (.ab1) files and a wild-type reference file are uploaded to the INDIGO webserver using Selenium WebDriver and ChromeDriver. Next, **Step 3: Analysis**, For each.ab1 file, the program performs the following sub-steps. (a) **Extract Page Source**: The program extracts relevant data (e.g., DNA sequence data) from the.ab1 file. (b) **Highlight PAM Sequence**: It identifies and highlights Protospacer Adjacent Motif (PAM) sequences. The program repeats this analysis for the next.ab1 file in the folder until all files are processed. In **Step 4: Retrieve Results**, the pipeline downloads the HTML reports for further review. Then, **Step 5: Cleanup**, clears temporary files or browser sessions. In **Step 6: Results**, the pipeline organizes the processed results, looping back to process the next.ab1 file in the folder (“Repeat for next. ab1 file”). This cycle continues until all files are processed, and the program ends (“End Program”).

### Performance Evaluation

The pipeline was tested on.ab1 files generated from Sanger sequencing of CRISPR-Cas9 edited rice lines. These files covered two guide RNAs (gRNAs) designed to create specific mutations in the target region of rice genomic DNA. In the Script we have provided a highlighting region which can be used to identify the PAM containing region of gRNA sequences (color coded) taken from rice target gene (Figs. 2 and 3). Each file was analyzed using a matching wild-type reference.ab1 file, and the pipeline successfully produced HTML reports with PAM regions highlighted in each sample (Fig. 4). The figures depict the sequence alignment outcomes derived from the INDIGO result page interface, comparing deconvoluted sample sequences with wild-type sequences. Two guide RNAs, gRNA-1 and gRNA-2, were analyzed, with indels highlighted: blue for gRNA-1 and yellow for gRNA-2. In alignment 1, the gRNA-1 sequence matches with the wild-type, marked by blue, whereas the gRNA-2 deconvoluted sequence lacks yellow highlighting due to the deletion. In alignment 2, both gRNA-1 and gRNA-2 sequences match with wild-type sequence, indicating that there are no mutations in allele 2. These results offer valuable insights into the genetic alterations induced by the dual gRNA system, emphasizing the varying effects of indels.

**Fig. 2:**
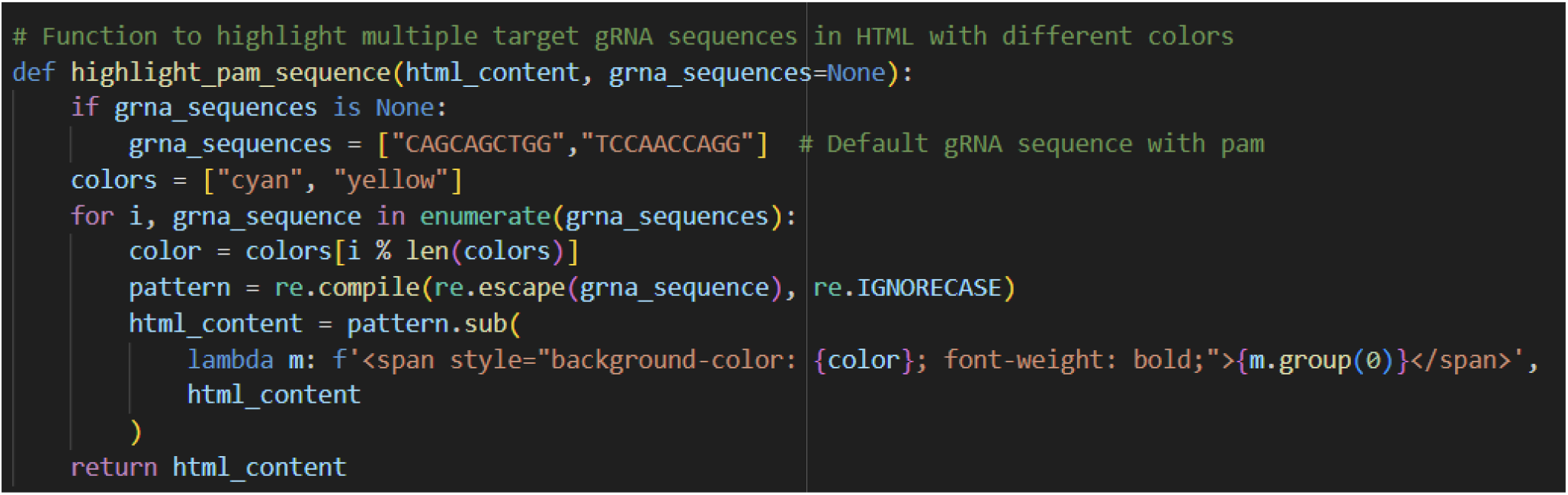
Script for highlighting PAM containing regions. The sequence information is taken from rice target gene.

**Fig. 3:**
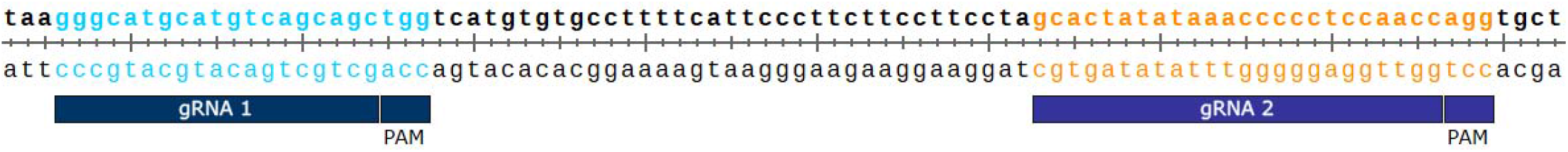
Targeted gRNA region’s along with PAM sequence from SnapGene. gRNA 1 is highlighted in blue and gRNA 2 is highlighted in yellow.

**Fig. 4:**
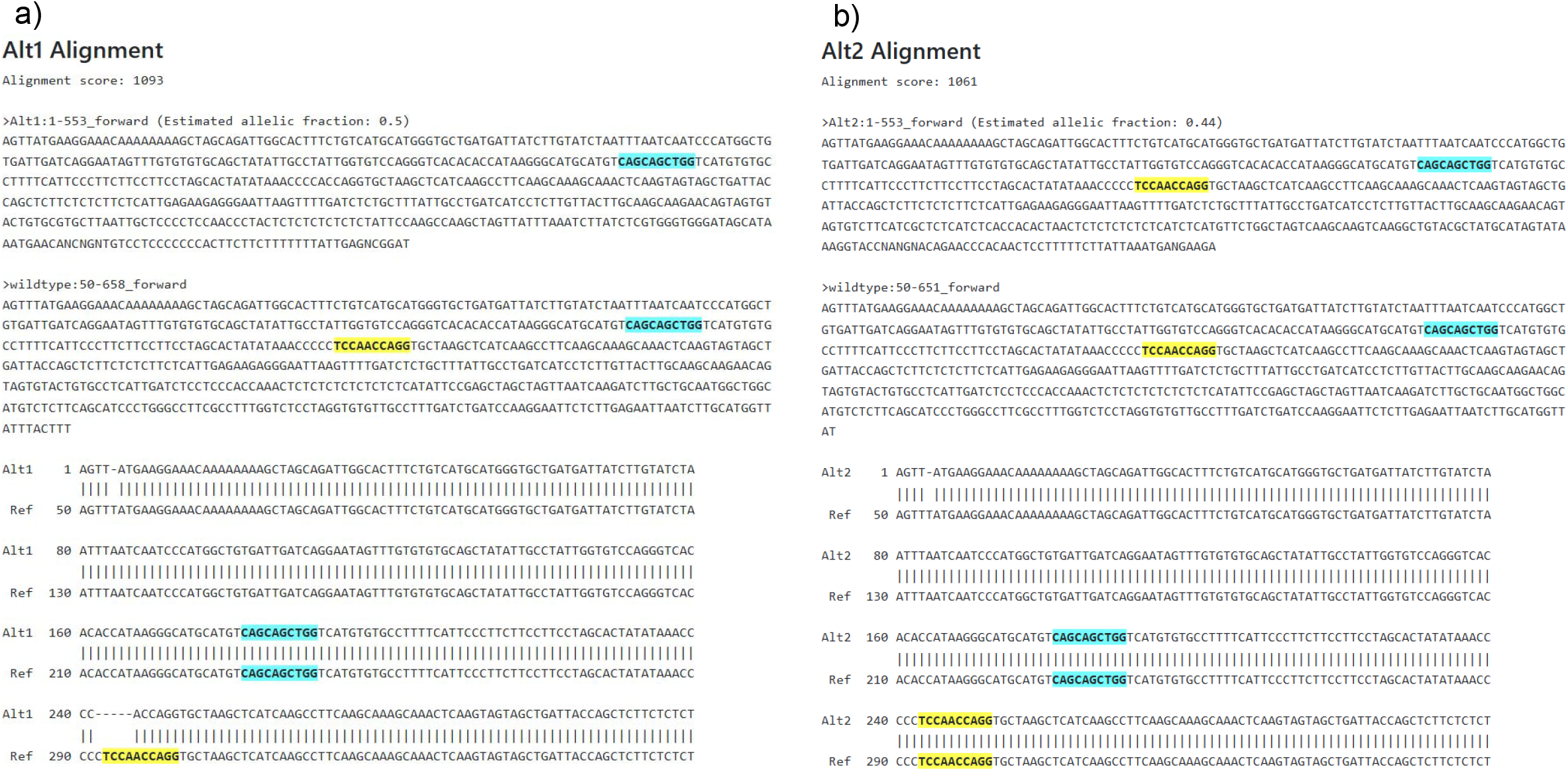
INDIGO result page showing the deconvolution of .ab1 files followed by alignment of sequencing reads against a reference sequence (wild-type). Annotated DNA sequence showing two guide RNA target regions, each highlighted in a different color (blue and yellow) to distinguish their positions within the sequence. a) Alt1 alignment (allele1 1 alignment), b) Alt2 alignment (allele 2 alignment).

### Accuracy

The pipeline demonstrated high accuracy in detecting indel mutations. When compared to manual analysis, the automated results matched 100% in identifying indels and their positions relative to the PAM sequence. The PAM regions highlighted in the HTML reports were accurate in all cases, as confirmed by SnapGene Viewer annotations (Fig. 3). The use of INDIGO’s reliable alignment and variant detection algorithms further ensured precise mutation identification across sample data (Fig. 4).

### Efficiency

The automation pipeline significantly improved the efficiency of mutation analysis, compared to manual methods. Manual analysis of.ab1 files would have been time-consuming, requiring researchers to individually upload files, review results, and annotate PAM regions. In contrast, the automated pipeline completes the same task in limited time on a Windows 10 system. The command prompt interface displays the file upload confirmations and processing status for each input file and the time clearly indicates the efficiency of the automation developed to process large numbers of Sanger sequencing files (Fig. 5). Its ability to handle multiple gRNA regions in a single run further enhanced efficiency, eliminating the need for repeated manual interventions.

**Fig. 5:**
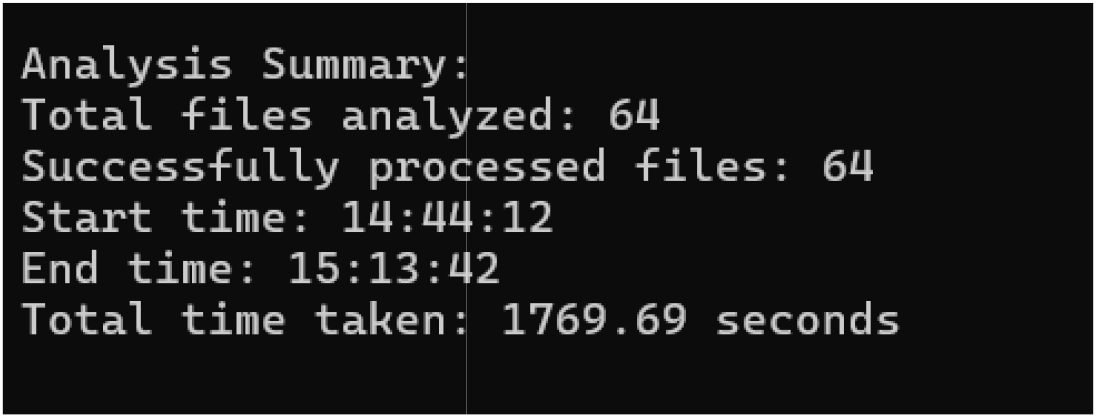
Screenshot of the command prompt interface of automated sequence analysis pipeline. The command prompt interface is used to execute the Python script for processing .ab1 files. The terminal displays the script’s output, including file upload confirmations and processing status for each input file and the time.

### Reliability

The pipeline proved highly reliable, successfully analyzing the files without issues. Its error handling mechanisms effectively managed problems like missing files or web element failures, either logging the errors or continuing with other files. In the rare cases of failure, caused by temporary network disruptions, the pipeline logged the errors and proceeded with the remaining files, ensuring overall workflow stability. Running the pipeline multiple times on the same dataset produced consistent HTML reports and PAM highlights, confirming its reproducibility and robustness for analysis.

Overall, the pipeline offers a reliable, efficient, and accurate solution for automating INDIGO webserver, making it well-suited for high throughput analysis of Sanger sequence files in plant genome editing research.

## Discussion

This automated pipeline streamlines CRISPR-Cas9 mutation analysis by processing Sanger sequencing data via the INDIGO webserver. Utilizing Selenium for web automation and incorporating PAM sequence highlighting, it boosts efficiency, scalability, and interpretability for high-throughput plant genome editing. Future enhancements could include support for diverse sequencing file formats, integration with additional mutation analysis tools, and parallel processing to minimize runtime. This work highlights automation’s potential to revolutionize bioinformatics workflows, freeing researchers to focus on biological insights rather than repetitive tasks.

## Availability

The automation code is publicly available at https://github.com/TIGS-rice/Indigo_Automation

## Acknowledgements

We express our gratitude to the EMBL Genome Facility, for providing access to the INDIGO, Gear Genomics web application, which was instrumental in our data analysis. We also acknowledge the institutional support from TIGS, which facilitated this research and to all the lab members for their collaborative efforts and contributions.

## References

1. Jinek, M., Chylinski, K., Fonfara, I., Hauer, M., Doudna, J. A., & Charpentier, E. (2012). A programmable dual-RNA-guided DNA endonuclease in adaptive bacterial immunity. Science (New York, N.Y.), 337(6096), 816–821.

2. Ricroch, A., Clairand, P., & Harwood, W. (2017). Use of CRISPR systems in plant genome editing: Toward new opportunities in agriculture. Emerging Topics in Life Sciences, 1(2), 169–182. 10.1042/ETLS20170085

3. Feng, Z., Mao, Y., Xu, N., Zhang, B., Wei, P., Yang, D. L., Wang Z., Zhang Z., Zheng R., Yang L., Zeng L., Liu X. & Zhu, J. K. (2014). Multigeneration analysis reveals the inheritance, specificity, and patterns of CRISPR/Cas-induced gene modifications in Arabidopsis. Proceedings of the National Academy of Sciences, 111(12), 4632–4637. 10.1073/pnas.1400822111

4. Sanger, F., Nicklen, S., & Coulson, A. R. (1977). DNA sequencing with chain-terminating inhibitors. Proceedings of the National Academy of Sciences, 74(12), 5463– 5467. 10.1073/pnas.74.12.5463

5. Yang, H., Wu, J. J., Tang, T., Liu, K. D., & Dai, C. (2017). CRISPR/Cas9-mediated genome editing efficiently creates specific mutations at multiple loci using one sgRNA in Brassica napus. Scientific Reports, 7(1), 7489. DOI: 10.1038/s41598-017-07871-9

6. Rausch, T., Fritz, M.H., Untergasser, A. & Benes, V. (2020). Tracy: basecalling, alignment, assembly and deconvolution of sanger chromatogram trace files. BMC Genomics 21, 230. 10.1186/s12864-020-6635-8

